# Time-dependent effect of fluoride on caries lesions development in a rat caries model

**DOI:** 10.64898/2026.08.18.745531

**Authors:** Aditya Banerjee, Samhita Sunkara, Leticia Capalbo, Nicolas Yoshino, Livia M. A. Tenuta

## Abstract

Since model dose-response is critical when assessing caries lesion development over time, this study evaluated the influence of fluoride dose and treatment duration on caries progression in a rat caries model. *Streptococcus mutans*-infected Sprague-Dawley rats were treated with deionized water, 226 ppm F^−^, or 2,260 ppm F^−^ twice daily for 3, 4, or 5 weeks. Caries lesions were assessed using Larson’s modification of the Keyes scoring system and complemented by micro-computed tomography (µCxT). Intraoral fluoride availability, serum and bone fluoride concentrations and microbial counts were also determined. Fluoride reduced caries severity in a dose- and time-dependent manner. While early enamel lesions were detected in all groups, extensive dentine lesions increased over time, in a dose-dependent manner, in the control and 226 ppm F^−^ groups, and were not observed in the 2,260 ppm F^−^ group after 5 weeks. Intraoral and bone fluoride availability increased significantly with fluoride concentration and treatment duration, whereas serum fluoride levels reflected fluoride dose instead of treatment duration. µCT-derived enamel volume correlated negatively with both total and extensive caries scores, supporting its utility as an objective measure of lesion severity. In conclusion, extending model length from 3 to 5 weeks increased the severity of caries lesions in a dose-dependent manner. Fluoride intraoral availability and bone fluoride also demonstrated a dose and time-dependent response.

## Introduction

Rat caries models have been instrumental in elucidating the anticaries effect and mode of action of different therapies [1–3]. They enable precise control of diet, microbial challenge, treatment exposure, and sampling time, allowing integrated assessment of caries development, oral microbiology, and systemic safety. Importantly, such models permit direct assessment of disease endpoints through standardized caries scoring system [4], which remain a critical benchmark for evaluating anticaries efficacy across experimental interventions. Recent methodological advances such as the use of digital photographs and microcomputed tomography to help with caries scoring can further streamline the use of these models in the future [5]. Yet, as with any other model to study caries, it is important that appropriate dose-response to the factors under study is demonstrated. That is particularly critical in an *in vivo* caries model in which lesions initiate and progress in severity under exposure to anticaries treatments.

Previous studies have demonstrated that increasing fluoride concentration enhances caries protection in rats [6–8]; however, both insufficient exposure duration and excessive fluoride dosing can obscure biologically relevant relationships between fluoride availability, tissue incorporation, and caries outcomes. The importance of an adequate time frame to observe significant and relevant differences in anticaries agents has been previously discussed by early model developers [9]. Establishing a model with dose-response to fluoride requires adjusting cariogenic challenge, fluoride exposure and time of sample collection to ensure that lesions have progressed significantly and that tooth destruction has not become extensive as to obscure differences among groups.

In the present study, we investigated the time-dependent anticaries effect of fluoride treatments in a rat model over exposure periods of 3, 4, and 5 weeks, including intraoral and systemic fluoride measurements.

## Methods

### Experimental design

Sprague-Dawley rats (n=18) were randomized by litter to three treatment groups: deionized water (negative control), 0.05% sodium fluoride (226 ppm F^−^) and 0.5% sodium fluoride (2260 ppm F^−^). Treatments were performed twice/day and animals euthanized after 3, 4 and 5 weeks (n=2 animals, one male and one female, per group, per time). Variables analyzed were intraoral fluoride bioavailability after treatments, microbial counts and caries score, as well as fluoride concentration in the serum and femur.

### Rat caries model

Two litters of 15 days-old Sprague-Dawley rat pups (9 males and 9 females) were obtained with their dams. Negative prior infection with *Streptococcus mutan*s was confirmed by an oral swab and inoculation on mitis salivarius bacitracin (MSB) agar (0.2 U/mL bacitracin, supplemented with 15% (w/v) sucrose) when animals were 18 days-old. Animals were infected on two consecutive days with 0.4 mL of a suspension of *S. mutans* UA159 in saline solution supplemented with 1% sucrose, via oral swabbing. Infection was confirmed on the following day. Animals were fed a 56% sucrose diet (NIH Diet 2000, TD.171001.PWD, Teklad, Envigo) throughout the experiment. After weaning, the animals were housed separated by groups and sex, in ventilated wire-bottom cages.

Treatments (100 µL) were applied twice/day, in two increments of 50 µL delivered to each side of the mouth using a pipette. Weights of the animals were measured weekly. After 3, 4 or 5 weeks, animals were euthanized by CO_2_ asphyxiation, followed by blood collection through cardiac puncture. Animals were decapitated, and the left mandible was aseptically dissected and immersed in 5 mL of sterile saline, for microbial counts; the right femur was dissected for bone fluoride assessment. Both mandibles and the maxilla were defleshed after heating the heads in a pressure cooker for 10 min.

### Intraoral fluoride availability

Animals scheduled to be euthanized after 5 weeks were used. At 4 and 5 weeks, 5 min after treatment application, a soft brush, previously washed with 0.5 M HCl and rinsed abundantly with deionized water to remove any fluoride contamination, was rolled in the animal’s mouth, on both sides, and the content transferred to 50 uL of deionized water. Fluoride concentration was determined in a fluoride electrode adapted for microanalysis [10, 11], after neutralization with TISAB III, and expressed as ng of fluoride collected.

### Microbial counts

The left mandible was aseptically collected into 5 mL of sterile 0.9% (w/v) NaCl in a 15 mL tube. Tubes were sonicated for 2 min (10 s on, 10 s off, 20% amplitude) in a cup horn (QSonica) coupled to a sonicator (Fisherbrand, Thermo Scientific). Serial ten-fold dilutions were prepared and plated in duplicate on Todd Hewitt (TH) agar, following incubation at 5% CO_2_, 37°C, for 24-48 h. Colony morphology was used to identify mutans streptococci, expressed as the log_10_ of CFU per mandible.

### Caries lesions estimation

Caries lesions detection and scoring were performed using Larson’s modification of the Keyes system [4]. Smooth-surface caries, including buccal, lingual, and morsal surfaces, were examined visually under a stereomicroscope, and confirmed after staining with 0.03% murexide [12]. For sulcal and proximal lesion assessment, molars were sectioned longitudinally in the mesiodistal plane through the center of the crown. To capture overall caries burden, the earliest detectable lesion score, indicating enamel involvement, was computed across all evaluated surfaces and is referred to hereafter as the “all lesions score”. Advanced dentine involvement was also evaluated as an indicator of extensive caries lesions by using the Dx score [4].

In addition to the conventional scoring, mandibles from 11 animals were scanned before sectioning using micro-computed tomography (µCT, Scanco Medical AG, Switzerland) and the micrographs were then segmented using the Dragonfly software [13]. Firstly, median filtering was applied to the raw images following which they were segmented using the ‘Universal Jaw Segmentation_V2023_1’ model in Dragonfly. Using the brush tool, multislice painting was performed throughout the slices to ensure well-defined segmentation of enamel, dentin and bone. Mineral density and voxel volume of the enamel was calculated by using the ‘Bone Mineral Density’ feature in Dragonfly. Volume of both mandibles enamel was summed up and expressed as mm^3^.

### Systemic fluoride concentration

Blood collected through cardiac puncture was centrifuged (7,500 x g, 4°C) and the serum collected. Serum fluoride concentration was determined using the hexamethyldisiloxane (HMDS) microdiffusion method coupled with a fluoride ion-selective electrode (Orion 96-09, Thermofisher Scientific, USA) [14]. Fluoride concentration was measured potentiometrically using the fluoride ion-selective electrode calibrated with standards processed identically by microdiffusion and expressed in micromolar.

For bone fluoride concentration determination, femurs were cleaned of adherent soft tissue, rinsed with deionized water, and air-dried. A 4 mm-long section was cut from the middle portion.

The outer and inner diameters of the bone section were measured using a caliper, and the inner and outer surface area of exposed bone were calculated and summed. Bone sections were added to 0.4 mL of 0.5 M HCl at room temperature in an orbital mixer at 200 rpm to extract surface-accessible fluoride. After 1 min, the acid extract was immediately neutralized with an equal volume of TISAB II containing 0.5 M NaOH. Fluoride concentration in the neutralized extract was measured using the fluoride ion-selective electrode calibrated with fluoride standards prepared with the same reagents as the samples. Fluoride values were expressed as µg F^−^/cm^2^ of exposed bone surface.

### Statistical analysis

Data of each variable were analyzed using two-way ANOVA, considering the factors treatment and time. Data which did not fit the assumptions of ANOVA was transformed. The correlation between the caries scores and enamel volume obtained from microCT assessment was investigated for “all lesions” and “extensive lesions” using Pearson coefficient of correlation. A significance level of α=5% was used. Data was analyzed using GraphPad Prism for Mac, version 11.0.

## Results

### Caries lesions develop early, but severity increases over time in a dose-dependent manner

When assessing all lesions, no increment in caries score within each treatment group was observed from 3 to 5 weeks, with the highest fluoride concentration group showing significantly lower scores than the other groups (Table; Figure 1). On the other hand, considering only extensive dentine lesions, an increment over time was observed for the negative control and the lower fluoride concentration group (the highest fluoride concentration group did not present extensive lesions). Also, extensive lesions differentiated the treatment groups in a dose-dependent matter, an effect which seems to be better observed after 5 weeks.

**Figure 1.**
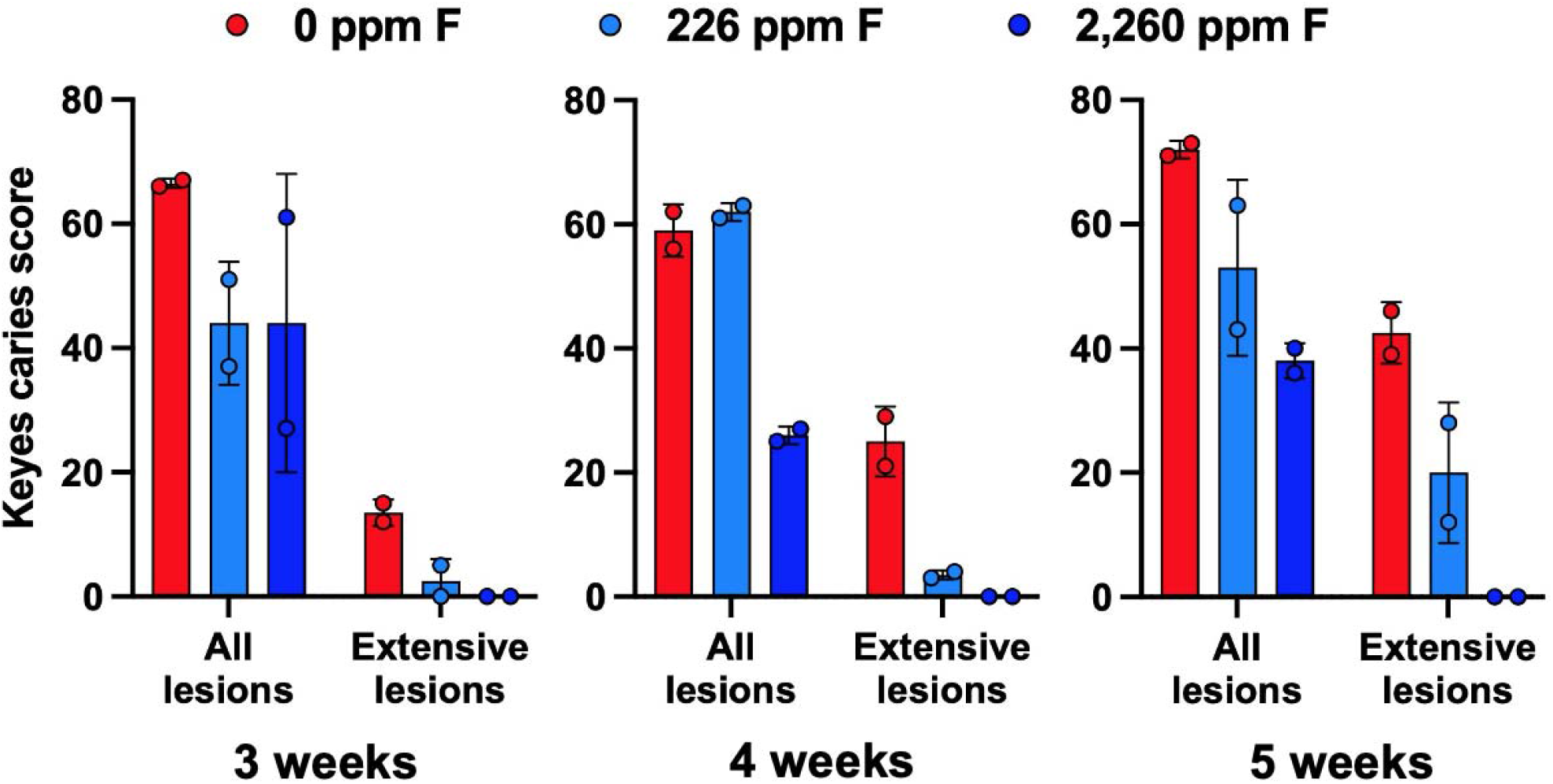
Caries lesions quantified using Larson’s modification of the Keyes system in rats treated with deionized water (0 ppm F), 226 ppm F and 2,260 ppm F for 3, 4 and 5 weeks. All lesions include any caries lesion, starting at enamel lesions. Individual values of two independent biological replicates (n = 2) are shown; bars represent the average and error bars indicate the standard deviation. For statistical comparisons, please refer to Table 1.

#### Higher doses of fluoride increase intraoral fluoride bioavailability

Intraoral fluoride levels increased in a dose- and time-dependent manner. Compared with the control group, rats receiving 226 ppm F showed elevation in intraoral fluoride bioavailability, with increases of approximately 1.2- and 15.7-folds at 4 and 5 weeks respectively, indicating sustained enrichment of fluoride within the oral environment during continuous exposure (Table). This effect was markedly greater in the 2,260 ppm F group, in which intraoral fluoride content increased by 93.5- and 470.2-folds across the same time points.

**Table:** Effect of fluoride concentrations and time (3, 4 or 5 weeks) on the variables (average ± SD, n=2).

| Variables | Weeks | 0 ppm F |  | 226 ppm F |  | 2,260 ppm F |  |
| --- | --- | --- | --- | --- | --- | --- | --- |
| **Intraoral fluoride after treatment (ng F)** | 4 | 0.34 ± 0.14 | Aa | 0.44 ± 0.06 | Aa | 33.7 ± 12.7 | Ab |
| | 5 | 0.39 ± 0.04 | Aa | 5.70 ± 3.53 | Bb | 169.7 ± 57.2 | Bc |
| **Microbial counts (log<sub>10</sub>CFU/mandible)** | 3 | 7.7 ± 0.08 | | 7.7 ± 0.06 | | 7.9 ± 0.04 | |
| | 4 | 7.9 ± 0.21 | NS | 8.2 ± 0.19 | NS | 7.6 ± 0.06 | NS |
| | 5 | 7.8 ± 0.16 | | 7.8 ± 0.11 | | 7.5 ± 0.41 | |
| **All lesions (Keyes caries score)** | 3 | 67 ± 0.7 | | 44 ± 9.9 | | 44 ± 24.0 | |
| | 4 | 59 ± 4.2 | a | 62 ± 1.4 | a | 26 ± 1.4 | b |
| | 5 | 72 ± 1.4 | | 53 ± 14.1 | | 38 ± 2.8 | |
| **Dx lesions (Keyes caries score)** | 3 | 14 ± 2.1 | Aa | 2.5 ± 3.5 | Aab | 0.0 ± 0.0 | Ab |
| | 4 | 25 ± 5.7 | Aa | 3.5 ± 0.7 | Ab | 0.0 ± 0.0 | Ab |
| | 5 | 43 ± 4.9 | Ba | 20 ± 11.3 | Bb | 0.0 ± 0.0 | Ac |
| **Serum fluoride (μM)** | 3 | 0.09 ± 0.016 | | 0.10 ± 0.020 | | 0.11 ± 0.024 | |
| | 4 | 0.09 ± 0.013 | a | 0.09 ± 0.003 | a | 0.14 ± NA | b |
| | 5 | 0.08 ± 0.007 | | 0.10 ± 0.016 | | 0.14 ± 0.009 | |
| **Bone fluoride (μg/cm<sup>2</sup>)** | 3 A | 0.03 ± 0.003 | | 0.28 ± 0.032 | | 1.82 ± 0.44 | |
| | 4 AB | 0.04 ± 0.002 | a | 0.27 ± 0.038 | b | 2.27 ± 0.21 | c |
| | 5 B | 0.05 ± 0.003 | | 0.31 ± 0.002 | | 2.58 ± 0.08 | |

Data was analyzed by two-way ANOVA, for the effects of treatment, time and their interaction. Data followed by the same letter are not statistically different ( $p > 0.05$ ). Statistically significant interactions are represented by upper case (for the comparison among time points) and lower case (for the comparison among treatments) letters. When only the main effects were significant, the effect of treatment is shown by lower case letters after each fluoride treatment; when only the effect of time was significant, the effect is shown by upper case letters after the weeks. Data of intraoral fluoride after treatment and bone fluoride were transformed to log<sub>10</sub> to fit the assumption of ANOVA. NS = not significant. NA = not available (only one sample was available for analysis, so no standard deviation could be calculated).

### Higher fluoride doses increase its concentration in serum and accumulation in bone

Twice daily treatment with a 2,260 ppm F solution significantly increased circulating fluoride in serum when compared with the other groups, with no significant effect of experiment time (Table). On the other hand, bone fluoride concentration increased significantly over time. Compared to the deionized water-treated animals, bone fluoride increased by about 7.5-folds in the 226 ppm F group and 60-folds in the 2,260 ppm F group (Table).

### Mandible molar enamel volume correlates with caries scores

By segmenting data of enamel from both mandible molars, and obtaining the enamel volume, we were able to observe a negative and significant correlation between both caries score metrics presented here (all lesions and extensive lesions) and enamel volume (Figure 2).

**Figure 2.**
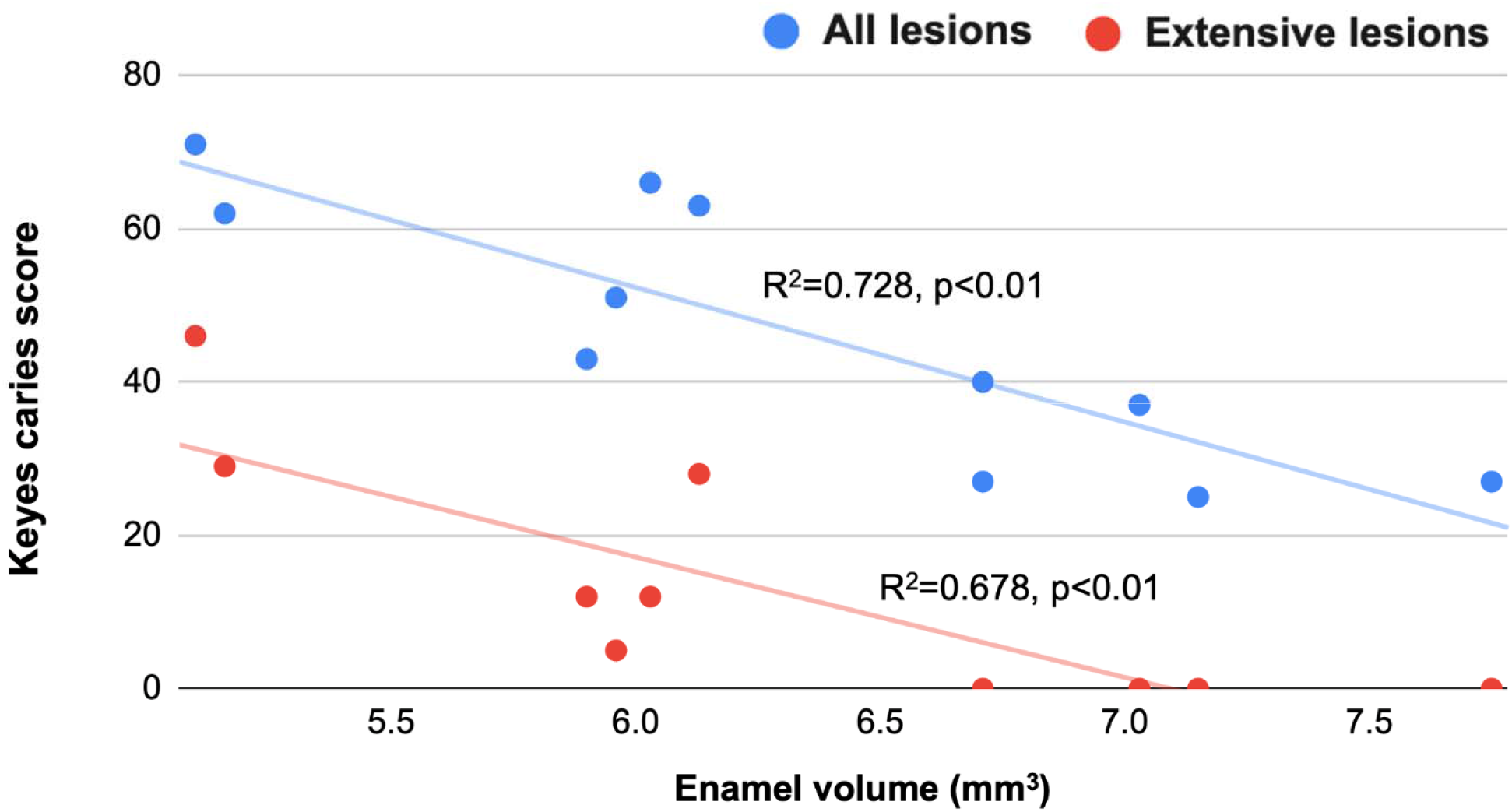
Correlation between Keyes caries scores (all lesions or extensive lesions) and mandible enamel volume obtained from microCT analysis.

## Discussion

The present study demonstrates that fluoride exerts a strong dose- and time-dependent protective effect against dental caries in a rat model, mainly shown by a reduction in lesion severity over time. Although lesions formed in all groups early in the assessment period (3 weeks), their severity was significantly impaired by a higher fluoride concentration; after 5 weeks, no animals in the highest fluoride group exhibited extensive lesions. Our results also demonstrate that time has a significant effect on intraoral fluoride bioavailability fluoride accumulation in mineralized tissues. By integrating caries quantification and systemic fluoride measurements, our findings provide a comprehensive view of how temporal exposure to varying fluoride doses collectively regulate the development of caries in a rat model.

Our data show that systemic fluoride accumulation is dose-dependent, with the greatest increases observed in bone compared with serum. This is consistent with the well-established pharmacokinetics of fluoride, in which circulating fluoride reflects recent exposure, while bone serves as the primary long-term reservoir due to fluoride’s affinity for mineralized tissue and its incorporation into hydroxyapatite during remodeling [15]. Notably, by the time of the euthanasia, which happened at least 15 h after the last treatment, only the highest fluoride groups showed slightly elevated serum fluoride levels. In contrast, bone fluoride content was substantially elevated at both fluoride doses, confirming that skeletal incorporation is highly sensitive to daily fluoride exposure [15, 16].

To the best of our knowledge, this seems to be the first study investigating intraoral fluoride availability after a topical treatment in this type of model. We found that increasing fluoride doses significantly increase intraoral fluoride availability over time. This observation supports the concept that although the applied treatment affects intraoral fluoride concentration, the possibility of retaining fluoride in intraoral reservoirs may also play a role [17–19]. The striking increase in detected fluoride between 4 and 5 weeks suggests an increase in intraoral fluoride retention sites over time. That could be the result of an increase in the size of the animals (from 4 to 5 weeks, males body weight increased from 186.1±14.8 g to 203.3±3.1 g, and females from 153.2±10.1 g to 180.8±14.5 g), and therefore the intraoral surface area, as the oral mucosa has been shown to be an important intraoral retention site [20]. It is also possible that caries progression might offer additional retention sites, with cavities offering an increase in tooth surface area and also in biofilm accumulation. The method used to estimate intraoral fluoride in this study, rolling a brush inside the animal’s mouth, would not allow a differentiation to be made among these different fluoride sources, but it seems to be an additional way of investigating the anticaries effects of therapies applied intraorally. Although in this study no antimicrobial effect of fluoride could be observed, assessing intraoral availability of different therapies could allow for the plausibility of additional anticaries properties, such as antimicrobial effects, to be determined. For example, inhibiting fluoride export by oral microbes can make them more susceptible to fluoride are lower concentrations [21], enhancing the antimicrobial effect of fluoride [22, 23].

A strength of this study is the combined use of Keyes scoring and µCT-based quantification of enamel loss. Traditional lesion scoring is semi-quantitative and subject to the interpretation of the examiner. The inclusion of µCT imaging allowed objective measurement of mineral density and volumetric enamel loss, thus correlating with lesion severity. This integration of lesion scoring with µCT supports previous work demonstrating that µCT can complement conventional caries scoring by enabling volumetric mineral loss assessment and improving reproducibility in rat-caries models [5, 24].

The time-dependent nature of fluoride effects observed here is also important. Caries progression in rat models is influenced by the balance between cariogenic challenge and protective factors, and differences in fluoride efficacy may not be detectable if lesions are either too early to diverge or too advanced such that severe cavitation masks protective effects. The present findings align with earlier observations that an adequate experimental duration is required to establish robust dose-response relationships and to distinguish between partial and maximal anticaries effects [9]. Our data suggest that prolonged exposure, within the timeframe tested here, not only increases fluoride availability but also increases the likelihood of detecting differences in lesion severity and mineral loss across groups.

Overall, our findings support the classical understanding that fluoride acts primarily through sustained topical effects within the intraoral niche, reducing the speed of tooth demineralization. They also emphasize that both dose and exposure duration are critical determinants of measurable anticaries efficacy in rat models.

## Acknowledgements

The authors would like to acknowledge the insightful discussions about microCT data analysis with Dr. Tomer Stern, University of Michigan School of Dentistry, Dr. Lucia Cevidanes, The University of North Carolina at Chapel Hill Adams School of Dentistry, and Dr. Murat Maga, University of Washington Department of Pediatrics.

## Statement of Ethics

This study protocol was approved by the Institutional Animal Care and Use Committee, University of Michigan [# PRO00011647].

## Author contributions

CRediT: **A. Banerjee:** Investigation, Writing – original draft; **S. Sunkara:** Investigation, Writing – review and editing; **L. Capalbo:** Investigation, Writing – review and editing; **N. Yoshino:** Investigation, Writing – review and editing; **L.M.A. Tenuta:** Conceptualization, Investigation, Formal analysis, Funding acquisition, Writing – original draft, Writing – review and editing. All authors reviewed the final version of the manuscript and agree to be accountable for all aspects of the work.

## Data Availability Statement

All data generated in this study is compiled in the manuscript’s table and figures. Additional details, like individual values for each variable, can be requested from the corresponding author.

## Use of generative artificial intelligence

No generative artificial intelligence has been used to create or review text in this manuscript.

